# Frequency-Selective Responses to Temporal Interference Stimulation in Hippocampal Neurons

**DOI:** 10.64898/2026.11.24.768895

**Authors:** Raul de Palma Aristides, Leonardo L. Gollo, Francesca Castaldo, Giulio Ruffini, Santiago Canals, Claudio R. Mirasso

## Abstract

Temporal interference (TI) stimulation aims to engage deep brain structures noninvasively by superimposing two slightly detuned high-frequency electric fields (e.g. ¿ 2 kHz), producing a low-frequency envelope (1–120 Hz) where they overlap. Existing accounts explain why a neuron responds to the envelope, but none identifies what determines which beat frequency a given neuron amplifies most. We address this using morphologically detailed, conductance-based models of hippocampal CA1 pyramidal neurons and four interneuron subtypes, varying carrier frequency, envelope frequency, field amplitude, and orientation. Since the applied field contains no spectral component at the beat frequency, that response must first be generated by nonlinear membrane dynamics; we characterize the frequency-dependent gain applied to it once generated. That gain is band-pass: slow voltage-gated currents suppress the response at low frequencies and produce a peak at the resonant frequency. Modulated stimulation therefore outperforms an unmodulated carrier only when the envelope frequency matches the resonance of the relevant compartment, near its firing threshold. Notably, this advantage extends to carriers as low as 250 Hz, well below the minimum reported for simplified neuron models (1400–1700 Hz). Under distributed synaptic drive the advantage is lost, although frequency selectivity persists. Selectivity also varies across targets: because resonance depends on conductance composition, the optimal envelope frequency differs across compartments and cell types, and augmenting resonant conductances in parvalbumin-positive basket cells confers an advantage they otherwise lack. Carrier frequency is therefore less constrained than commonly assumed, and envelope frequency should be matched to the resonant properties of the target neuron type.

## 1 Introduction

Temporal interference (TI) stimulation has emerged as a promising non-invasive approach for selectively activating specific neural populations while minimizing stimulation of surrounding tissue [1]. The technique enables the targeting of deep brain structures by superimposing two high-frequency electric fields (*>*1kHz) with a small frequency offset (1-120Hz), generating a low-frequency envelope capable of modulating neuronal activity at the interference focus [2]. TI stimulation has been validated in humans, where focal targeting of the hippocampus was shown to modulate hippocampal activity and enhance the accuracy of episodic memories in healthy participants [3]. This spatial selectivity has generated considerable clinical interest, with potential applications ranging from deep brain stimulation of the globus pallidus internus and the subthalamic nucleus for Parkinson’s disease [4, 5] to the modulation of hippocampal circuits in epilepsy [6], bipolar disorder [7], and depression [8].

Despite this enormous potential, the biophysical mechanisms by which neurons respond to TI stimulation remain incompletely understood. Grossman and coworkers [2] attributed TI efficacy to the low-pass filtering properties of the neural membrane, which attenuate high-frequency carrier oscillations while transmitting the slow envelope to the membrane potential. Subsequent computational work has argued that passive filtering alone cannot account for the effect, since a linear filter preserves the frequency content of its input, and that an ion-channel mediated rectification process is required [9]. A related account identifies a cycle-by-cycle balance between inward and outward currents that holds for an unmodulated carrier and is disrupted when the envelope changes rapidly; in the squid Hodgkin-Huxley model under current injection, this mechanism operates above a carrier of ≈ 1700 Hz, which was linked to the sodium activation time constant remaining below 0.6 ms [10, 11]. While existing accounts address why a neuron responds to the envelope at all, none of them identifies which envelope frequency a given neuron will amplify most.

This gap widens in biophysically detailed neurons, which have been reported not to exhibit TI stimulation in isolation, contrary to experimental results [11]. Either the single-cell substrate depends on features that simplified models omit, such as resonance-generating conductances distributed over an extended dendritic arbor, or the effect is not a single-cell phenomenon at all and arises from recurrent excitation and inhibition. Three questions therefore remain open, all concerning the single neuron: (i) what biophysical mechanism underlies frequency-selective responses to TI stimulation in realistic neurons; (ii) what determines which envelope frequencies are most effective for a given neuron; and (iii) whether such a mechanism persists under ongoing synaptic drive. Answering these is a prerequisite for, not a substitute for, a network-level account of TI.

A candidate mechanism that could address these gaps is subthreshold membrane resonance: the tendency of neurons to respond preferentially to inputs at a specific frequency, arising from the interaction between passive membrane properties and active ionic currents such as the M-type potassium current (*I*_M_, mediated by Kv7 channels [12]) and the hyperpolarization-activated current (*I*_h_, mediated by HCN channels [13]) [14]. Resonance has been extensively characterized in hippocampal neurons [15, 16, 17] and is known to shape their selectivity for oscillatory inputs [18]. If the TI envelope acts as an oscillatory input, the neuron’s resonant frequency may influence which envelope frequencies are most effective, affecting frequency-selective TI responses tuned to each neuron’s intrinsic properties. Because the applied field contains no spectral component at the beat frequency, the response there must first be generated by nonlinear membrane dynamics; what the frequency-dependent impedance of the membrane then determines is how strongly that component is amplified.

Here we test this hypothesis using morphologically detailed, conductance-based models of hippocampal CA1 pyramidal cells and four interneuron subtypes: parvalbumin-positive basket cells (PVBCs), axo-axonic cells (AAs), Ivy cells, and oriens-lacunosum moleculare cells (OLMs), derived from the Blue Brain Project hippocampal circuit model [19, 20, 21]. We focused on the hippocampus for three reasons. First, hippocampal neurons are among the most characterized neuronal populations in the mammalian brain, with extensively mapped cell-type diversity and intrinsic electrophysiological properties [22, 23]. Second, it was the preparation in which TI stimulation was first characterized experimentally [2] and has since been among the most studied targets in TI studies [24], making it a relevant system against which to benchmark a mechanistic account. Finally, the hippocampus is a canonical model system for studying neuronal computation and neuromodulation: its well-characterized circuitry, diverse interneuron complement, and prominent role in learning, memory, and spatial cognition [25, 26] provide both the biological motivation and the cell-type diversity necessary to examine how resonance properties vary across a functionally defined neural circuit.

We establish this first in the simplest case, a neuron driven by injected current and an applied field with no synaptic input. Here the advantage of the modulated field is unambiguous: it appears when the envelope frequency matches the resonance of the target compartment, is gated by proximity to the spike threshold, and varies systematically across cell types according to their complement of active conductances, providing a biophysical basis for frequency-selective, cell-type-specific modulation. We then identify the limits of this account under more realistic conditions. Under distributed synaptic drive, the membrane fluctuates rather than settling near threshold, and the modulated field no longer outperforms an unmodulated carrier; what survives is the frequency selectivity itself, broadened and requiring larger field amplitudes to be detectable. Together, these findings establish intrinsic resonance as the single-cell substrate of frequency-selective TI sensitivity, and show that whether this substrate produces an advantage over an unmodulated carrier depends on the membrane’s operating point.

## 2 Results

### 2.1 Subthreshold resonance amplifies modulated field sensitivity near the spiking threshold

We first analyzed a morphologically detailed CA1 pyramidal neuron model, shown in Fig. 1a), obtained from the Blue Brain Project hippocampal circuit [19, 20, 21], before extending the study to different CA1 interneuron classes (see Methods 4.1). Given the nature of voltage-gated ionic channels, far from the spiking threshold, membrane conductances are effectively constant, and the neuron behaves approximately as a linear system. Under these conditions, oscillatory inputs, such as an oscillatory electric field, are filtered according to the membrane’s linear properties, with no selective response at the envelope frequency. Frequency sensitivity emerges only when voltage-dependent conductances are sufficiently recruited. Closer to the spiking threshold, the activation of these nonlinear currents endows the neuron with intrinsic resonance properties [14, 27, 28, 29].

**Figure 1:**
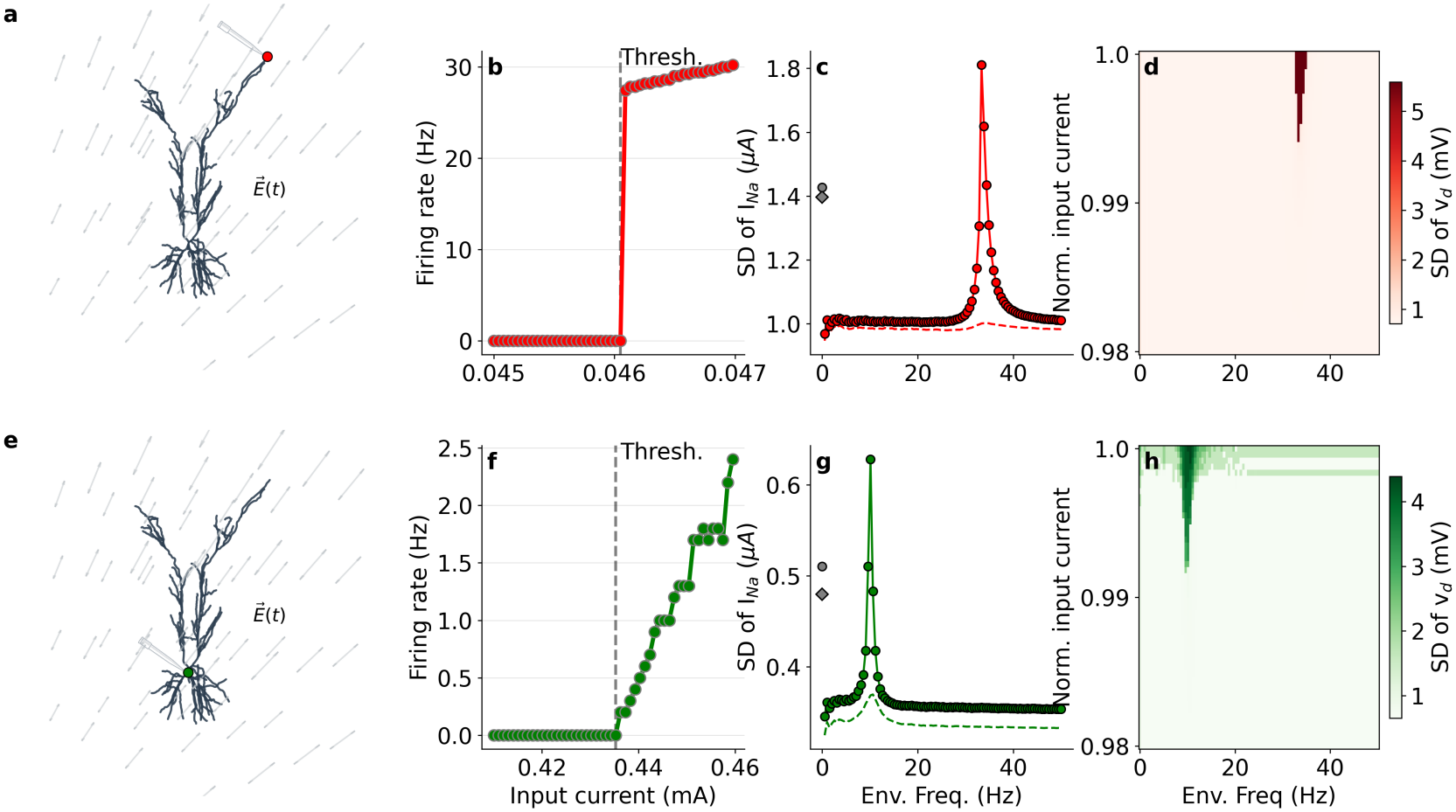
CA1 pyramidal neuron response to amplitude-modulated electric fields. a, CA1 pyramidal cell morphology, where red and green markers indicate the distal dendritic and somatic segments, respectively. At these segments, a steady current was injected (illustrated by the pipette) to control channel availability while the response to an oscillating electrical field, illustrated by the grey arrows, was measured. b, Firing rate as a function of injected current in the absence of an electric field. The neuron transitions abruptly from quiescence to sustained firing at a well-defined threshold current, defining the distance-to-threshold axis used in panels c-d. c, Response to an external electrical field as a function of envelope frequency, quantified by the standard deviation (SD) of the sodium current. Results are shown for a fixed carrier frequency of 2500 Hz and *E*_amp_ = 85 V/m. Curves are shown for two injected current levels: near firing threshold (*Ĩ* = 0.992) and far below threshold (*Ĩ* = 0.99). The resonant increase in *I*_*Na*_ at the envelope frequency emerges only near threshold, while the response far from threshold is largely insensitive to envelope frequency. d, Joint dependence of the response on injected current and envelope frequency. The heatmap shows the SD of the segment’s membrane voltage as a function of injected current (y-axis, up to the firing threshold identified in b) and envelope frequency (x-axis, 0-50 Hz), confirming that the resonant peak is confined to a narrow band near threshold across the full frequency sweep. e-g, as in b-d, but for the somatic segment.

Hence, we characterized the firing rate–input current (f–I) relationship of the dendritic compartment by injecting a steady current (marked as the red dot and the pipette in Fig. 1a). Fig. 1b shows a discontinuous transition from silence to firing in the dendritic segment. The current at which the neuron transitions from quiescence to sustained firing defines the threshold current *I*_thresh_. All subsequent results are reported as a function of the injected current normalized to this value,

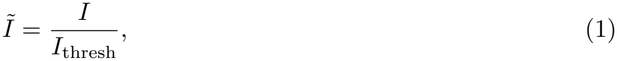

so that *Ĩ* = 1 corresponds to the threshold current and *Ĩ <* 1 to the subthreshold regime (see Methods 4.2).

Next, we exposed the neuron to a superposition of two extracellular sinusoidal fields with equal amplitude. When the two frequencies were identical, the superposition reduced to an unmodulated carrier; when they differed by Δ*f*, it produced an amplitude-modulated field with envelope frequency Δ*f* and maximal amplitude *E*_amp_ (see Methods 4.4). Keeping *E*_amp_ = 85 V/m, we swept the envelope frequency from 0 to 50 Hz at two steady input currents, one close to threshold and one far from it, and measured the standard deviation of the sodium current in the dendritic segment (Fig. 1c). Far from threshold, the unmodulated response (grey square) exceeded the modulated response at every envelope frequency. Closer to threshold, this ordering reversed over a narrow band, with the modulated response peaking at Δ*f* ≈33 Hz above the unmodulated reference (grey circle).

At this preferred frequency, the envelope acts as a selective drive, producing a larger sodium current response than the unmodulated alternative. The two-parameter space of input current versus envelope frequency (Fig. 1d) shows that this advantage is confined to a narrow band near threshold. Within it, the modulated field tuned to the resonant frequency cooperates with the subthreshold input to trigger action potentials before the nominal current-clamp threshold is reached, whereas the unmodulated field fails to elicit spiking. Finally, we repeated this paradigm at the soma, shown in panels Fig. 1e–g. While the transition from silence to spiking differs quantitatively between the dendritic and somatic segments (panels b and e), the core phenomenon remains qualitatively the same: near the preferred envelope frequency, the modulated signal selectively amplifies local depolarization, lowering the threshold for spiking across both domains (panels d and h).

As Fig. 1c and f show, the soma and the distal dendrite resonate at different frequencies, which follows from how resonance arises. A slow regenerative current acts as an effective inductance, opposing the membrane’s passive capacitive relaxation and producing a preferred frequency at which impedance is maximal [14]. In hippocampal neurons, this role is played by currents such as *I*_h_ [13] and *I*_M_ [12]. Because these conductances are distributed non-uniformly along the somatodendritic axis, each compartment resonates at its own frequency. In this CA1 pyramidal cell, for instance, *I*_h_ conductance increases with distance from the soma along the apical dendrite, a well-established gradient [17, 30] that leaves the soma and the distal dendrite with different local channel compositions, effective inductances, and capacitive loads. Compartment-specific resonance frequencies along the dendritic trunk are discussed further in the Appendix (see Fig. 8).

To better understand the differential response to modulated inputs, Fig. 2 shows the transmembrane ionic currents for two envelope frequencies and the unmodulated condition at the somatic compartment, with *Ĩ* = 0.992. We focus on four representative currents: *I*_Na_, the total potassium current *I*_K_ (summing all potassium mechanisms in the model), T-type calcium (*I*_CaT_), and delayed rectifier potassium (*I*_Kdr_), each shown in a separate column. The first row shows the currents for when the envelope frequency Δ*f* matches the resonant frequency obtained in Fig. 1f. The second row shows the same comparison for Δ*f* = 40Hz. The third row shows the power at the envelope and carrier frequencies for both modulated conditions.

**Figure 2:**
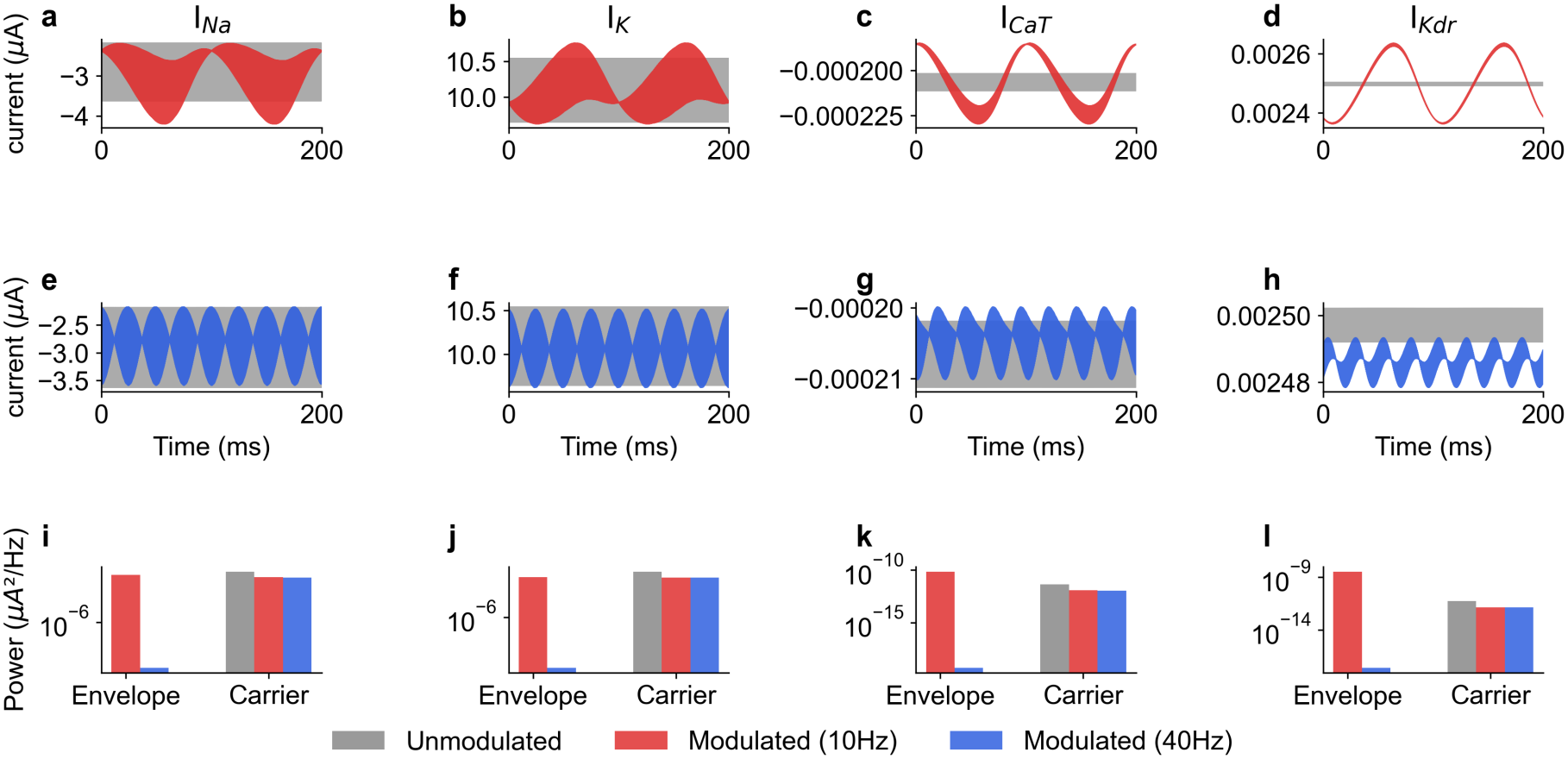
Time series of somatic transmembrane currents illustrate the distinct ionic responses to modulated and unmodulated stimulation. a-d, *I*_Na_, *I*_K_, *I*_CaT_, and *I*_Kdr_ (one current per column) in response to an unmodulated electric field (grey) and to a modulated field with envelope frequency matching the somatic resonant frequency (color), both at a carrier frequency of 2500 Hz, *E*_amp_ = 85V/m and *Ĩ* = 0.992. e-h, as in a-d, but for an envelope frequency that does not match the resonant frequency. i-l, Power spectra (*µA*^2^*/Hz*) of each current (columns as in a-h), computed at the carrier frequency for all three conditions (unmodulated, resonant, non-resonant) and at the envelope frequency for the two modulated conditions (envelope power is undefined for the unmodulated condition by construction). Stimulation at the resonant envelope frequency selectively increases power at the envelope frequency relative to the carrier, an effect that is absent or strongly reduced when the envelope frequency does not match resonance; the slow, high-threshold *I*_Kdr_ shows near-complete redistribution of power to the envelope frequency, consistent with its kinetics acting as a natural low-pass filter on the carrier.

At the resonant envelope frequency, all four ionic currents exhibit pronounced low-frequency modulation that is absent under unmodulated stimulation and substantially reduced at the non-resonant frequency. This effect is evident in both the time-domain traces and the spectral decomposition, where envelope-frequency power emerges strongly in the resonant condition while carrier-frequency power remains largely unchanged. Despite large differences in absolute amplitude, Na, K, CaT, and Kdr currents all show enhanced envelope components near resonance, indicating that envelope extraction is distributed across multiple conductances rather than arising from a single channel type. For instance, the Na current shows 57% more power in the envelope than in the carrier, and the CaT current 60% more. The total potassium current shows only a 3% difference, since its components respond at different phases and their envelope contributions largely cancel in the sum. The delayed rectifier, taken alone, shows 2502% more power at the envelope than at the carrier.

Among the currents examined, *I*_Kdr_ shows the most spectrally selective response, with almost all of its power at the beat frequency. Slow kinetics alone do not confer this property: a current must be too slow to follow the carrier yet fast enough to follow the envelope, and *I*_Kdr_ falls in that band. Its high activation threshold (+13 mV) and slow kinetics preclude meaningful activation at kHz timescales, leaving negligible power at the carrier frequency. This selectivity is frequency-dependent: the envelope-to-carrier power ratio is large at the resonant envelope frequency (10 Hz) and substantially smaller at 40 Hz (Fig. 2l), so it reflects the channel’s kinetics rather than an artifact of near-zero carrier power. *I*_Kdr_ is nonetheless small in absolute terms (≈2.5 nA against ≈3 *µ*A for *I*_Na_, Fig. 2d versus 2a), so the ratio describes its spectral purity rather than a large contribution to the membrane response. As a voltage-gated conductance, it follows the slow component of the membrane potential regardless of which currents generate that component, so its selectivity characterises its response rather than its role in demodulation.

### 2.2 Boundaries of the modulated advantage in stimulus parameter space

We next characterize how the somatic response depends jointly on field amplitude and proximity to threshold, and whether this dependence is sensitive to the carrier frequency. Fig. 3 shows the difference in somatic firing rate between modulated and unmodulated stimulation as a function of field amplitude (x-axis, logarithmic) and proximity to threshold (y-axis), for six carrier frequencies: 250, 500, 1000, 2500, 5000 and 9000 Hz. This range covers frequencies below and above the sodium channel activation cutoff frequency reported for the squid Hodgkin-Huxley model under current injection [10, 11].

**Figure 3:**
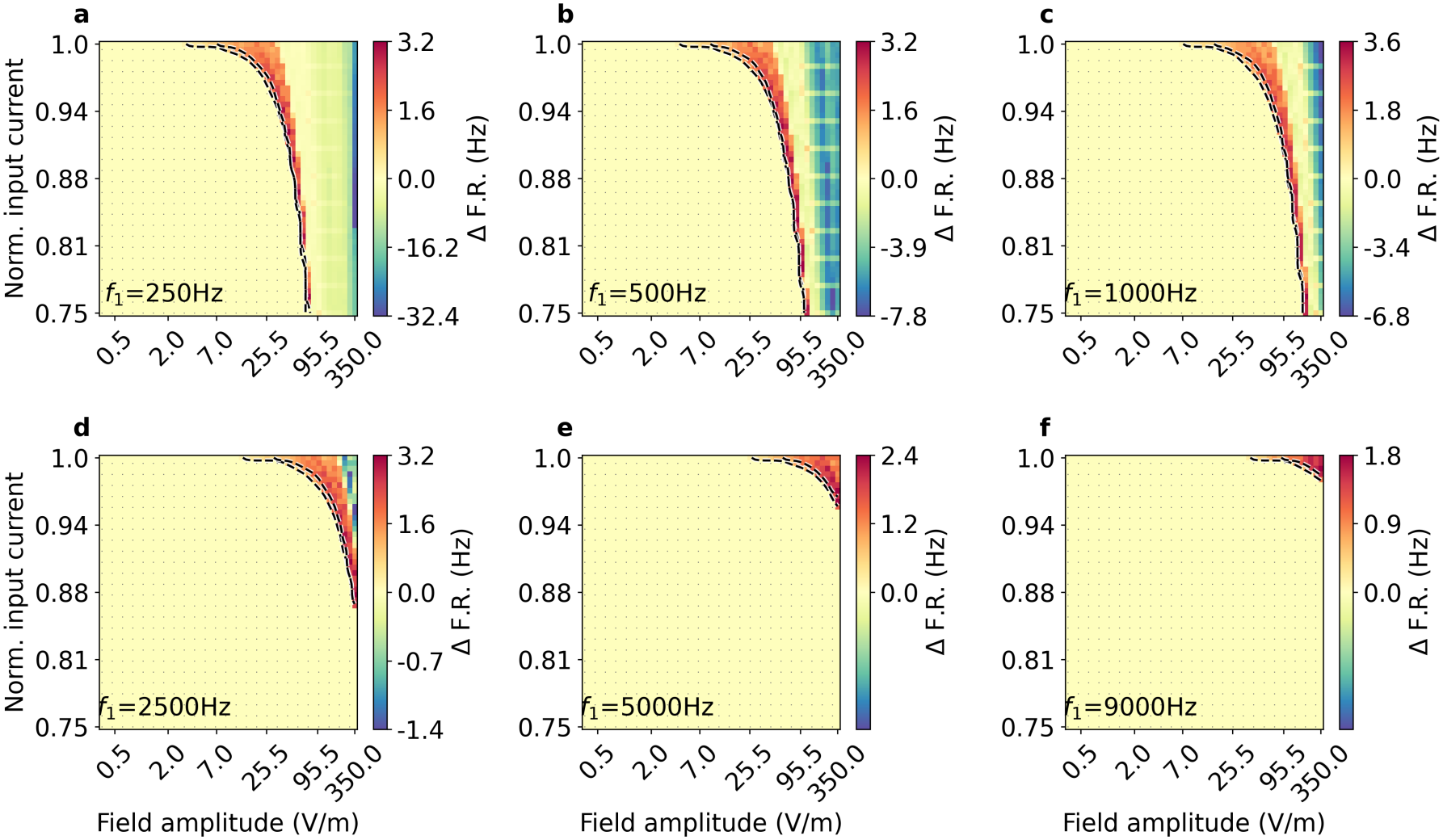
Input current–field amplitude response. Panels (a–f) show the response for carrier frequencies of 250, 500, 1000, 2500, 5000, and 9000 Hz. The color scale represents the firing rate difference, ΔF.R. = F.R.(*E*_mod_) − F.R.(*E*_unm_), where F.R.(*E*_mod_) is the firing rate evoked by an amplitudemodulated field whose envelope matches the neuron’s natural frequency, and F.R.(*E*_unm_) is the response to the corresponding unmodulated field, both measured at the soma. Positive values (red) indicate a modulated advantage, negative values (blue) an unmodulated advantage; the color scale is normalized independently for each panel and for each sign, with zero at the midpoint, so saturation is not comparable across panels. Dots mark where neither field elicited firing F.R.(*E*_mod_) = F.R.(*E*_unm_) = 0), which is otherwise indistinguishable from equal non-zero rates on the color scale. Dashed contours delimit the regions in which each condition elicits firing; the band between them is the parameter space in which the modulated field drives spiking while the unmodulated field does not. Contours are smoothed for display. The y-axis shows the normalized input current (0.75–1.0), with 1.0 corresponding to the minimum current required to elicit spiking (Fig. 1). The x-axis shows the electric field amplitude on a logarithmic scale, spanning 0.3–350 V/m.

Two observations emerge clearly from Fig. 3. First, firing requires both larger field amplitudes and closer proximity to threshold, confirming that channel availability gates the response as established in Fig. 1. Neither field elicited any response over much of the parameter space (dotted region), and this silent regime expands with carrier frequency. Absolute firing-rate maps for each condition are provided in the Appendix. Second, the modulated advantage is concentrated near threshold and at moderate field amplitudes, and is bounded: within the band delimited by the dashed contours the modulated field elicits firing where the unmodulated field does not, whereas at the highest amplitudes the unmodulated field produces the stronger response. Away from threshold, the advantage diminishes and increasingly stronger fields are required to produce a comparable effect. The two conditions are plotted separately rather than as a difference in the Appendix (see Figs. 14 and 15).

A modulated advantage is present at 250, 500, and 1000 Hz, below the threshold reported for the squid Hodgkin-Huxley model, which is therefore not a universal minimum carrier frequency. Its extent nonetheless diminishes as the carrier frequency increases, consistent with progressive attenuation by the membrane. The envelope frequency falls within the resonant bandwidth of the compartment, where membrane impedance is higher, which is what the kinetic account does not by itself predict. The field amplitude axis spans 0.3 to 350 V/m, covering the range from non-invasive human TI setups to in vitro slice preparations and mouse in vivo experiments [31, 32, 33].

Next, we examined how electric-field orientation affects the neuronal response, repeating the conditions from Fig 2: an unmodulated sinusoidal drive (2500 Hz carrier) and amplitude-modulated signals at Δ*f* = 10 and 40 Hz. Field orientation was sampled at 100 angles uniformly spaced over 2*π* (3.6^*°*^ resolution) in each plane. For an elongated cable-like neuron in a quasi-uniform field, membrane polarization is governed to leading order by the field component along the cell’s principal axis,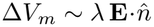, where *λ* is the electrotonic space constant [34, 35]. The response should therefore follow a cosine dependence on orientation, maximal when the field aligns with the somatodendritic axis and strongly attenuated when it is orthogonal [36, 37].

The morphology of the CA1 pyramidal cell is depicted across different planes in panels 4a–4c, while the response dependence on field orientation is illustrated in panels 4d–4f. In the *xy*-plane (Fig. 4a), the response is strongly orientation-dependent, peaking at alignment with the somatodendritic axis—where axial polarization between the apical dendrites and soma is maximized, leading to the greatest sodium channel recruitment. Away from this axis, the unmodulated input elicits a larger sodium influx than either modulated condition. The *xz*-plane (Fig. 4b) is transverse to the somatodendritic axis, resulting in a weak response dominated by the unmodulated input throughout. Finally, the *yz*-plane (Fig. 4c) contains the axis and demonstrates a pronounced orientation dependence, albeit at a reduced amplitude due to partial misalignment with the dominant dendritic direction.

**Figure 4:**
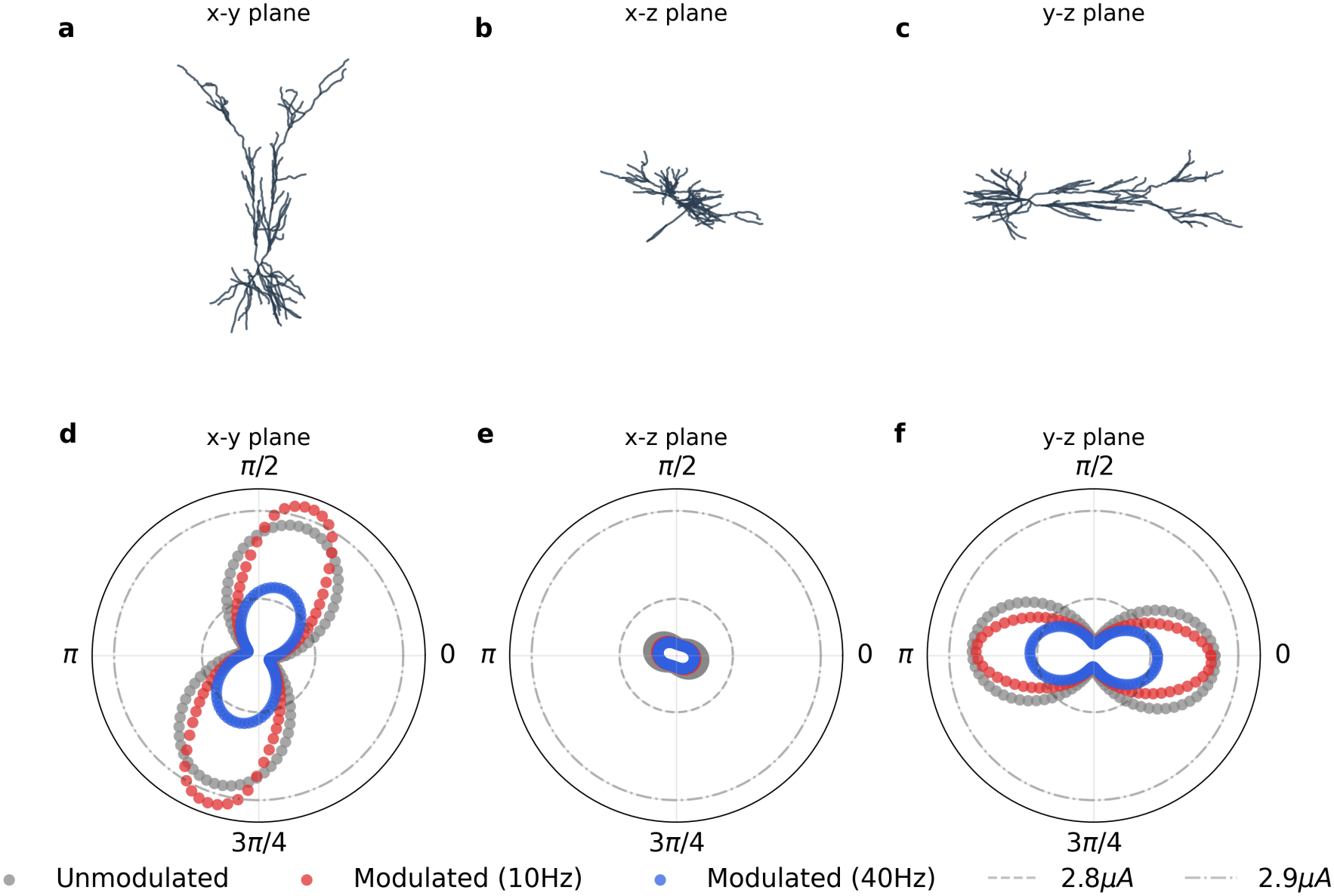
Effect of electric field orientation on the neuronal response. CA1 pyramidal cell morphology depicted across orthogonal projection planes in panels a–c (a: *xy*-plane, b: *xz*-plane, c: *y*-z plane). Polar plots showing the average sodium influx (*I*_*Na*_) as a function of field orientation in panels d–f for an unmodulated carrier (2500 Hz, gray) and amplitude-modulated signals at 10 Hz (red) and 40 Hz (blue) envelope frequencies. The response is maximal when the field aligns with the somatodendritic axis and scales with its projection onto that axis, consistent with the 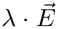 framework. The 10 Hz modulated input selectively outperforms the unmodulated one near the preferred orientation; away from this axis, and in planes with weak somatodendritic projection (*xz*), the unmodulated input dominates. Results obtained with *Ĩ* = 0.99 and .

### 2.3 Resonance under synaptic drive: broadened but preserved

Having established the response properties of CA1 pyramidal neurons under constant depolarizing drive, we next asked whether these effects persist in the presence of physiologically relevant synaptic input. This distinction is important because, in vivo, excitatory drive is primarily delivered through distributed dendritic synapses, rather than direct somatic or dendritic current injection, which can significantly alter neuronal dynamics [38, 39, 40]. We therefore replaced the constant current input with sustained dendritic excitatory synaptic bombardment (see Methods 4.5) and examined the firing-rate response to amplitude-modulated electric fields across different levels of synaptic conductance.

Under synaptic drive, the firing-rate response retained a frequency-dependent enhancement at sufficiently strong field amplitudes, as shown in Fig. 5. At *E*_amp_ = 300 V/m, amplitude-modulated stimulation produced a broad increase in firing rate centered around envelope frequencies of approximately 15–20 Hz, consistent with the resonant response observed under constant depolarization. The magnitude of this enhancement increased with synaptic conductance, reflecting the greater proximity of the membrane potential to spike threshold under stronger excitatory drive. Notably, the unmodulated stimulation condition (Δ*f* = 0 Hz) evoked a stronger firing-rate response than the amplitude-modulated inputs at both *E*_amp_ = 200 and *E*_amp_ = 300 V/m, indicating that continuous stimulation remained more effective than modulated waveforms in driving spiking activity. However, when the field amplitude was reduced to 200 V/m, the low-frequency enhancement was no longer detectable across the tested synaptic conditions. Thus, while synaptic excitation can amplify the firing-rate response once the resonance mechanism is engaged, the additional conductance load also raises the stimulation amplitude required to elicit this behavior. These results demonstrate that the resonance and associated firing-rate modulation observed under direct depolarization extend to a more physiological regime of dendritic synaptic bombardment.

**Figure 5:**
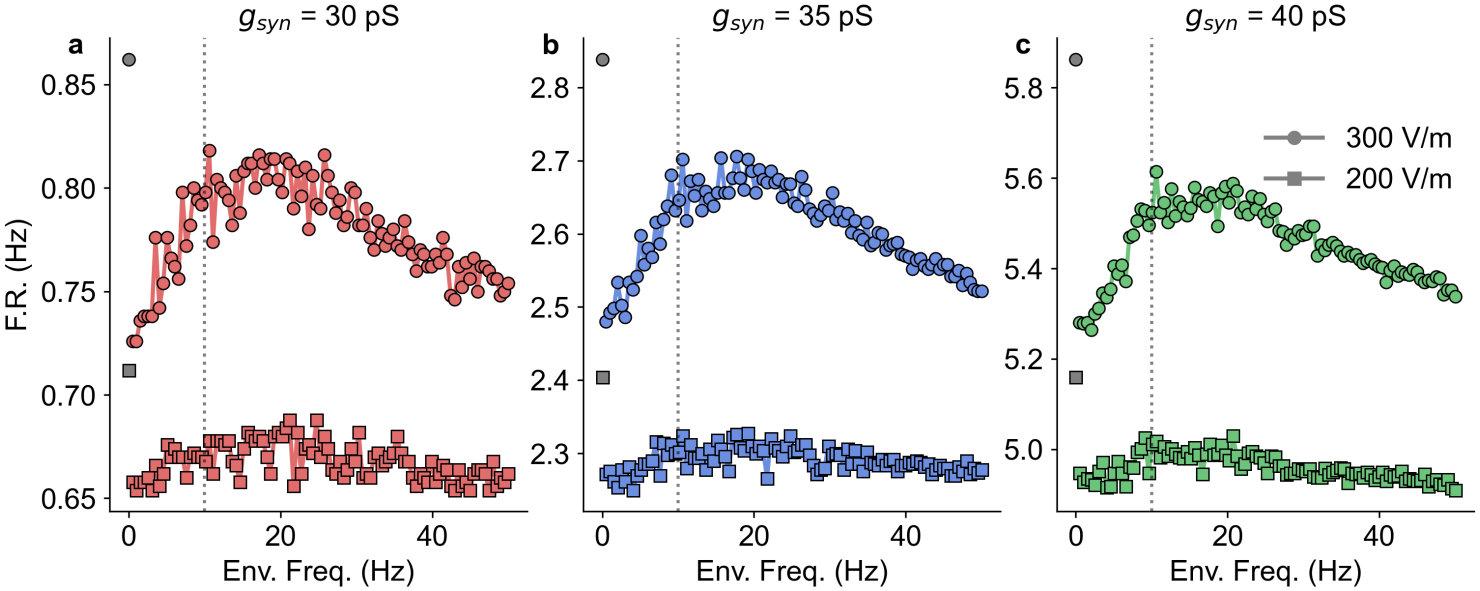
CA1 pyramidal neuron firing rate under synaptic bombardment and amplitude-modulated electric fields. Firing rate (FR) as a function of envelope frequency for two field amplitudes - *E*_amp_ = 300 V/m (circles) and *E*_amp_ = 200 V/m (squares) - and three synaptic conductance levels (a, 30 pS; b, 35 pS; c, 40 pS). The isolated grey markers represent the firing rate under unmodulated stimulation. NMDA-mediated synaptic input was distributed across the dendritic arbor at 50 Hz. At *E*_amp_ = 300 V/m, a broad resonant peak emerges around 10-20 Hz whose amplitude grows with synaptic conductance, consistent with stronger depolarization bringing the membrane closer to threshold. At 200 V/m, no resonant peak is detectable, indicating that synaptic conductance load raises the field amplitude required for the resonance mechanism to manifest.

### 2.4 Cell-type-specific resonance and its conductance dependence

Given the diversity of intrinsic electrophysiological properties among hippocampal interneurons, we examined how the response of the CA1 interneurons to amplitude-modulated electric fields. Figure 6 repeats the paradigm of Fig. 1 for OLM, PVBC, axo-axonic, and Ivy cells. The morphologies of each cell are illustraded in the first column, the resonant profile in the second and the the two-parameter space of input current versus envelope in the third. Both the clamp current and the response measurement are restricted to the soma, since these models lack active dendrites. Consistent with experimental impedance measurements in hippocampal neurons [18], we observed cell-type-specific differences in the response to oscillatory inputs. Interneurons differ both in their resonant frequencies and in their sensitivity to envelope frequency, supporting heterogeneous frequency-dependent filtering across interneuron classes.

**Figure 6:**
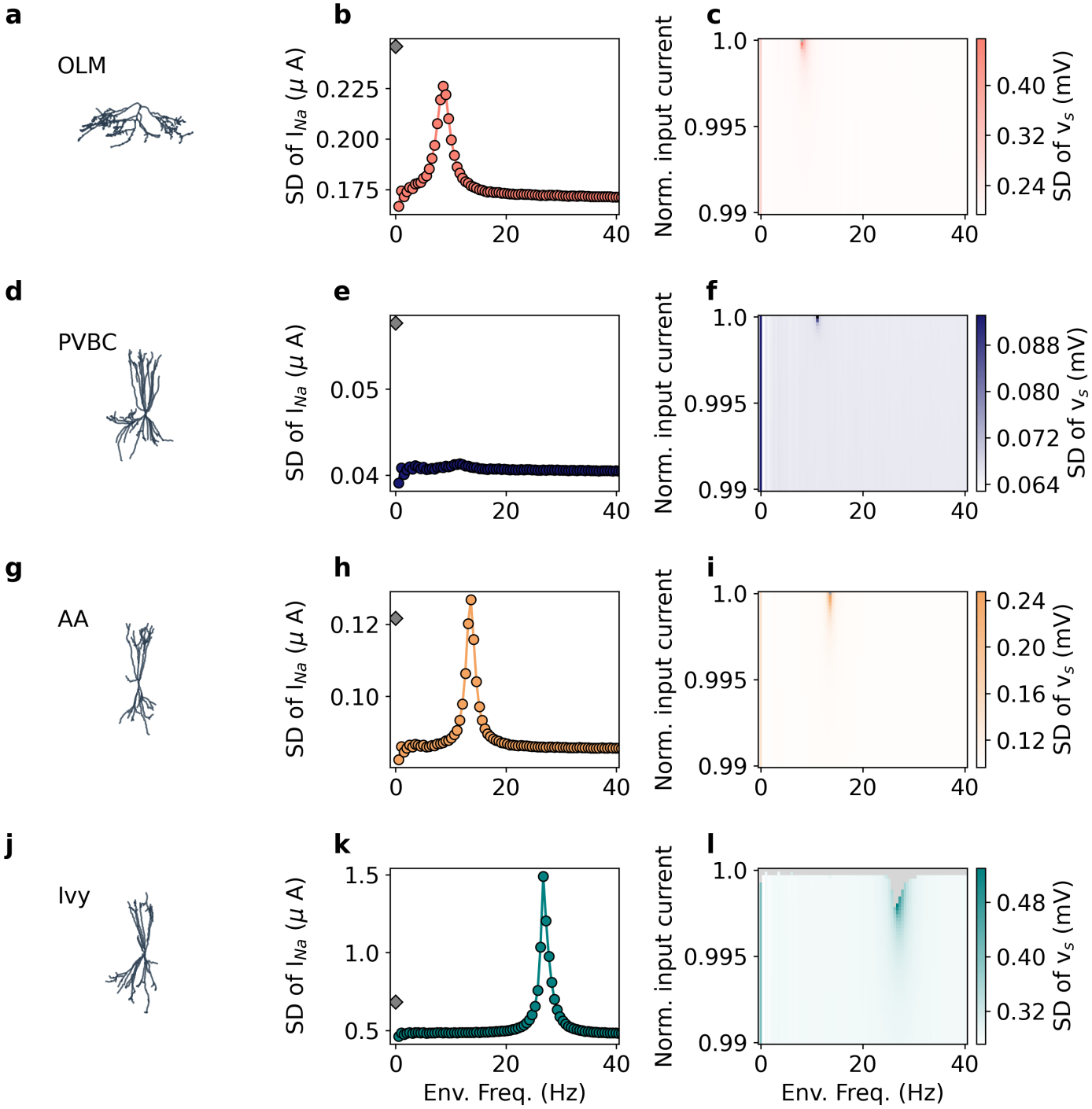
CA1 interneurons’ response to amplitude-modulated electric fields. a-c, OLM cell. a, Reconstructed morphology from the Blue Brain Project. b, SD of somatic *I*_*Na*_ current as a function of envelope frequency (with *Ĩ* = 0.995 2500 Hz carrier, *E*_amp_ = 85 V/m), showing a resonant peak that does not overcome the unmodulated response. c, Joint dependence on injected current and envelope frequency (colormap: SD of somatic membrane voltage), confirming that the resonant peak is confined to a narrow near-threshold band. d-f, as in a-c for the PVBC cell. g-i, Axo-axonic cell. j-l, Ivy cell. Each interneuron subtype exhibits a distinct resonant frequency determined by its active conductance composition, with the resonant advantage over unmodulated stimulation emerging only near threshold in all cases.

These differences are consistent with a biophysical origin rooted in intrinsic membrane conductances rather than passive properties alone. Variations in the expression of hyperpolarization-activated and other voltage-gated channels are expected to shape each cell type’s effective impedance profile, and thereby its preferred modulation frequency [14]. As in the CA1 pyramidal cell, this selectivity emerges within a restricted voltage range near spike threshold, consistent with a resonance mechanism engaged only under partially activated channel states [29]. The absence of a resonant peak in PVBC cells under TI stimulation does not indicate that the cell type lacks resonant properties. The same model shows a clear peak at 11 Hz when driven by a purely sinusoidal current (see Appendix), so the resonance is present but is not recruited by the amplitude-modulated field.

From the results in Fig. 6, we next asked whether the sensitivity of PVBC cells to AM electric fields can be enhanced by modulating intrinsic conductances implicated in subthreshold resonance. We focused on the hyperpolarization-activated current (*I*_h_) and the M-type potassium current (*I*_M_). As shown in Fig. 7a, increasing dendritic *I*_h_ conductance systematically strengthens neuronal resonance [17]. The maximal *I*_h_ conductance 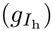 was scaled to 1.1×, 1.5×, 2×, 4× and 8×baseline. Across this range, the impedance profile becomes progressively more frequency-selective, with a monotonic increase in resonance amplitude, in accordance with previous studies [41].

**Figure 7:**
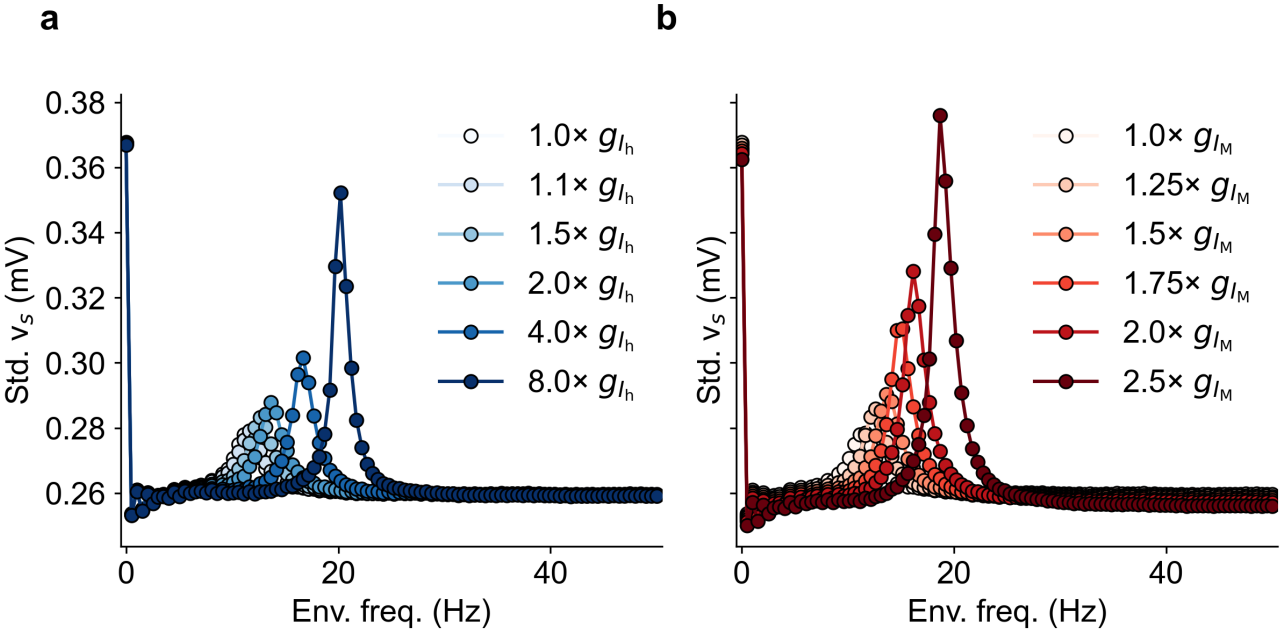
I_h_ and I_M_ currents modulate resonance and sensitivity to amplitude-modulated inputs in PVBC cells. a - Standard deviation of the somatic membrane potential (*v*_*s*_) as a function of envelope frequency for a 2500 Hz carrier and a fixed electric field amplitude of 280 V/m. Increasing the dendritic 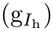 enhances the resonance peak and shifts the resonance frequency to higher values. b - Same as (a), varying the somatic and axonal 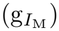. Increasing 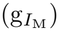 similarly enhances the resonance peak and increases the resonance frequency, but with substantially greater sensitivity, as a (∼ 2.5)-fold increase is sufficient to produce a comparable effect.

**Figure 8:**
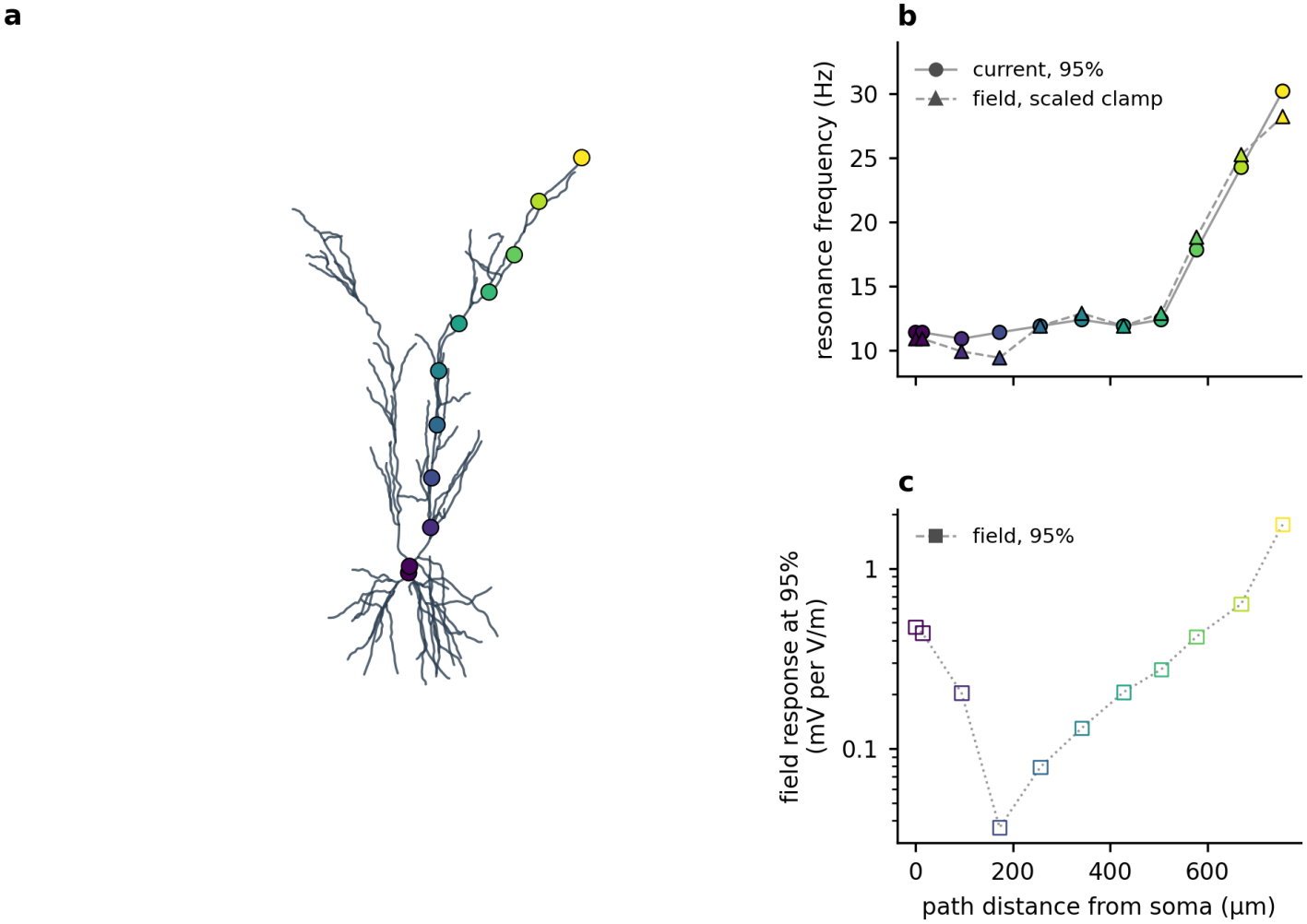
(a) CA1 pyramidal cell morphology (axon not shown) with the 11 stimulated locations, coloured by path distance from the soma; the same colours mark each location in (b) and (c). Each location was held by a DC current at a fraction of its own spiking threshold and driven sinusoidally at 1–50 Hz. (b) Resonance frequency (peak of the membrane-potential response at the drive frequency) vs path distance, for two protocols: local sinusoidal current at 95% of threshold (filled circles, solid line) and uniform field with the holding current scaled per location between 90% and 99.5% (filled triangles, dashed line). (c) Peak field-induced response per V/m at 95% of threshold, on a logarithmic scale; it is smallest in the mid-trunk.

Mechanistically, this arises because *I*_h_ acts as a voltage-dependent restorative conductance recruited during hyperpolarization. Increasing 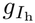 enhances this restorative feedback, improving suppression of slow membrane fluctuations and thus reducing low-frequency impedance. At the same time, high-frequency responses remain primarily constrained by membrane capacitance, yielding a more pronounced band-pass profile and a larger resonance peak. Overall, the impedance transitions from weak frequency dependence at baseline conductance to strong resonance at elevated 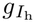 .

Increasing *I*_M_, a slow-activating voltage-dependent K^+^ outward current, produced a similar effect, as shown in Fig. 7b. Largely localized to the somatic compartment, *I*_M_ acts as a key resonant current by filtering low-frequency inputs and shaping subthreshold theta resonance in conjunction with *I*_*NaP*_ [12, 42, 43, 44, 45, 46, 47]. The effect of increasing 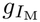 differs from that of increasing 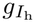 in two respects. First, the response to the unmodulated field decays with increasing 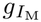 . This does not reflect a shift in operating point, since the injected current was re-normalised to the rheobase of each modified model, but rather the dependence of the field-evoked response on the conductance itself. Second, at 2.5*×* baseline conductance, the modulated response surpasses the unmodulated one. Augmenting a somatic resonant conductance is therefore sufficient to confer on PVBC cells the modulated advantage they otherwise lack. Both manipulations also shifted the resonant peak toward higher frequencies, consistent with the shorter effective membrane time constant produced by the added conductance.

## 3 Discussion

Resonance describes the ability of a neuron, or of individual segments within it, to respond selectively to inputs at a preferred frequency, set by the interaction between passive membrane properties and slow voltage-gated currents [14]. Our findings show how it shapes the response of CA1 neuronal models to TI stimulation. When the beat frequency generated by the two electric fields (*E*_1_ and *E*_2_) matches the resonance frequency of the stimulated compartment, responses are selectively amplified, and TI stimulation elicits action potentials even where the corresponding unmodulated high-frequency fields remain individually subthreshold (Fig. 1). When the beat frequency deviates from resonance, this amplification is lost, and the unmodulated carrier can produce a stronger neuronal response than the amplitude-modulated stimulus. Because resonance arises from voltage-gated conductances that are distributed non-uniformly along the somatodendritic axis, distinct compartments resonate at distinct frequencies, rising from ≈10 Hz at the soma and through the proximal apical trunk to ≈30 Hz in the distal tuft (Appendix Fig. 8). This provides a potential mechanism for frequency-selective modulation of distinct neuronal compartments.

### 3.1 Generation and amplification are separable stages

A neuron’s response to TI can be separated into two stages: the generation of a response at the beat frequency, and its subsequent amplification. Generation is necessary because the applied field contains no spectral component at the beat frequency, so a nonlinearity must create one. Luff et al. [48] showed that neurons perform this step, producing a difference-frequency component that is abolished by TTX and therefore originates in voltage-gated sodium channels, and Cao et al. [10] described the same step as the disruption of a cycle-by-cycle current balance. Neither account predicts which beat frequency is most effective, and the reason is that generation is largely insensitive to it: Luff et al. report comparable difference-frequency amplitudes across carriers spanning two orders of magnitude at a fixed 10 Hz beat frequency. What determines efficacy once that component exists is amplification by the membrane’s frequency-dependent gain, which is what our results characterise.

Our conductance manipulations separate the two stages directly. Raising somatic 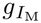 in PVBC cells amplified the beat-frequency component of the somatic voltage several-fold near the cell’s resonant frequency, left it unchanged at beat frequencies well away from the peak, and did not alter the carrier component (Appendix Fig. 5). Improved generation would have scaled all beat frequencies comparably and might have altered the carrier response; frequency-selective enhancement with an unchanged carrier is instead the signature of increased gain. The manipulation therefore acts on amplification, and the two stages can be separated experimentally.

### 3.2 Determinants of efficacy: amplitude, carrier, orientation and cell type

Where the unmodulated field prevails, it prevails on delivered energy rather than on waveform. Matching the two conditions by peak amplitude, as in the original experimental demonstration [2] and in subsequent modelling [10, 11], leaves the unmodulated field with twice the mean-square amplitude, since the cross term of the two detuned components averages to zero. Equalising mean-square amplitude instead reverses the ordering: the modulated response then exceeds the unmodulated reference at resonance and converges onto it away from resonance (Appendix Fig. 12). This does not make peak matching the wrong comparison. Grossman et al. applied all conditions at matched total current and found that two 100 *µ*A channels detuned by 10 Hz drove firing while a single 200 *µ*A channel at the same carrier did not, so the modulated field prevailed there despite the same energy disadvantage. The two matchings hold different stimulus quantities fixed and answer different questions: peak amplitude is the quantity constrained in practice, while mean-square amplitude isolates the contribution of the waveform itself.

The carrier range over which TI operates is wider than previously reported. A minimum carrier frequency of ≈1700 Hz has been derived for the squid Hodgkin-Huxley model under current injection, where it was linked to the sodium activation time constant [10, 11]. We find a modulated advantage well below this prediction (Fig. 3), so that figure does not generalise to morphologically realistic mammalian neurons. The practical consequence is that carrier selection is less constrained than currently assumed, which matters because higher carriers require larger field amplitudes to elicit comparable responses [49, 32], and the amplitude attainable non-invasively in humans is limited [50]. Field orientation relative to the somatodendritic axis is a further determinant of whether stimulation succeeds (Fig. 4), as anticipated from previous work [51, 37].

Cell type is the remaining determinant, and our findings converge with earlier work on this point. Cao et al. [10] reported that parvalbumin-positive interneuron models failed to exhibit TI stimulation where pyramidal models succeeded, and Caldas-Martinez et al. [11] found experimentally that 73% of layer 2–3 pyramidal cells but only 24% of PV cells exhibited TI responses. We likewise find PVBC cells to have the weakest resonance of the four CA1 interneuron subtypes examined, with no clear envelope-frequency preference (Fig. 6). These studies report the phenomenon without identifying its origin; our conductance manipulations supply one. The weak resonance of PVBC cells reflects their baseline conductance profile rather than a morphological constraint, since augmenting *I*_h_ or *I*_M_, both established contributors to resonance in hippocampal neurons [15, 16, 14, 52], confers the resonant advantage they otherwise lack (Fig. 7). Whether this relates to the fast feedback dynamics invoked by Caldas-Martinez et al. to explain the same cell-type difference remains to be established; the two accounts may well be complementary, since a cell with fast restorative dynamics and one with few slow resonant conductances would both show weak frequency selectivity.

Ivy cells were the only subtype in which the modulated field outperformed the unmodulated carrier, and they are among the most numerous interneuron classes in CA1, outnumbering the parvalbumin-expressing basket, bistratified and axo-axonic cells [53]. They are slow-spiking cells with dense fine axons innervating mostly the basal and oblique dendrites of pyramidal cells, to which they provide slowly rising and decaying inhibition. A stimulus that preferentially recruited this population would therefore act on pyramidal dendritic integration rather than on the perisomatic spike timing controlled by the parvalbumin-positive cells that showed the weakest response here. Whether this difference is exploitable depends on how the two populations interact under stimulation, which a single-cell study cannot determine.

### 3.3 Responses under synaptic drive

The same group also showed that pharmacological blockade of synaptic transmission strongly reduced TI-evoked responses, and concluded that TI stimulation is largely a network phenomenon [11]. Our results do not contradict that interpretation. They identify a complementary single-cell mechanism that becomes effective within a restricted perithreshold regime, where resonance-mediated amplification and active conductance dynamics become prominent, so that variability in membrane polarization, intrinsic excitability, and ion-channel composition may contribute to the heterogeneity of TI sensitivity across neurons. The models we used were optimized against electrophysiological features extracted from patch-clamp recordings in hippocampal slices, predominantly at the soma, with dendritic conductance distributions constrained by published gradients rather than fit directly [19, 20], and we simulated them in isolation by design. The field strengths required to elicit firing under current clamp (≈ 100 V/m) fall within the range reported in experimental TI studies in slice preparations and rodents [31, 32] but exceed those attainable in human studies [50].

Under distributed synaptic drive, the modulated field no longer outperformed the unmodulated carrier at any conductance tested, while frequency selectivity persisted in broadened form (Fig. 5). Random synaptic drive cannot reproduce the conditions the clamp imposes: the membrane fluctuates rather than settling near threshold, and stronger drive adds variance and conductance load instead of sustaining proximity to threshold [38, 39, 40]. The broadening follows from the same fluctuations, which move the membrane through a distribution of operating points each with a slightly different resonant frequency, so that the observed peak is an average over that distribution, widened in proportion to its variance; stochastic broadening of this kind is well established for resonant systems [54]. These are rate measures, following the criterion used in previous work, in which a neuron exhibits a TI response when the modulated field elicits firing where the unmodulated one does not [2, 11]; modulation could nonetheless organise spike timing within the beat cycle without changing the mean rate, and characterising this requires protocols designed for phase analysis, which we leave to future work. The negative result is informative precisely about where TI efficacy originates: if the advantage of modulation does not survive in an isolated neuron under realistic drive, while TI produces measurable effects in vivo [2, 3], then the effect is unlikely to be reducible to single-cell biophysics. What our results supply is the specification of what a network-level account would have to deliver, and the frequency dependence it would have to respect.

### 3.4 Limitations and future work

Two aspects of the present work lie outside what single-neuron simulations can address. The conditions under which neurons occupy the perithreshold regime in intact tissue are set by ongoing network activity, which we did not model; our results specify what a neuron does once it is in that regime, not how often it gets there. Recurrent interactions also modify the response itself, and work on transcranial alternating current stimulation indicates the direction, with active networks proving more sensitive to applied fields than isolated cells [55, 56, 57]. Frequency-selective responses are also observed at the circuit level: optogenetic drive of parvalbumin-positive interneurons at 40 Hz preferentially increases gamma power, while drive of pyramidal cells at 8 Hz increases theta power [58, 59, 60]. These preferences arise from recurrent excitation and inhibition rather than from intrinsic cellular resonance, and how the two interact is beyond the scope of a single-cell study. What the present results do establish is twofold: that the envelope frequency is a determinant of TI efficacy rather than a free parameter, with intrinsic membrane resonance setting that dependence, and that the carrier need not exceed the sodium activation cutoff.

A recent proposal places both the demodulation and its frequency tuning at the population level, with the firing-rate nonlinearity of a neural mass acting as the detector and the recurrent network as a resonant amplifier [61]. The two accounts are not exclusive: detection at the population level is inherited from the single neuron, and the cellular resonance characterised here would shape which frequencies the network receives.

### 3.5 Conclusions

Together, these findings suggest that intrinsic neuronal resonance provides a cellular substrate for frequency-selective sensitivity to amplitude-modulated electric fields, whose expression depends on the membrane’s operating point. The resonant advantage emerges within a narrow perithreshold regime, where voltage-dependent conductances are sufficiently engaged to sustain the nonlinear interaction. Under distributed synaptic drive, this frequency selectivity persists but becomes broader and more strongly dependent on field amplitude and the neuronal operating point.

## 4 Methods

### 4.1 Neuronal models

Neuronal morphologies were obtained from the Blue Brain Project (BBP) dataset via the Hippocampus Hub resource [19, 20, 21]. Computational simulations were carried out in the NEURON simulation environment [62, 63], with Python serving as the programming interface for model construction, simulation execution, and subsequent analyses. Detailed version information for Python and all software dependencies is provided in Table 1.

**Table 1:**
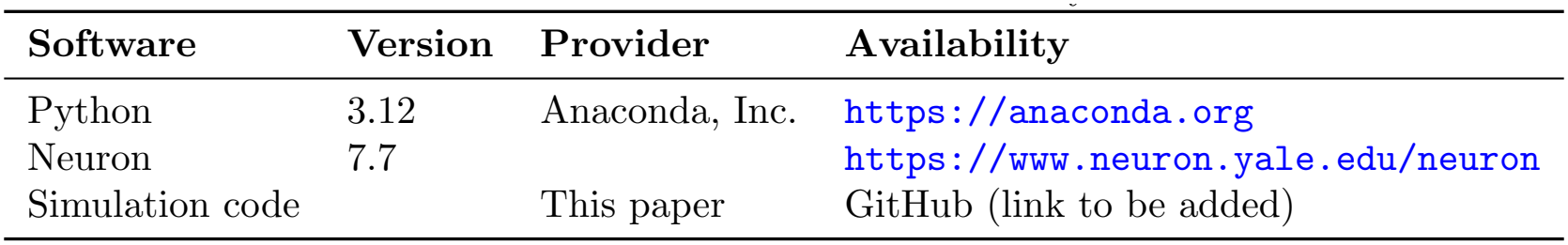
Software used in this study.

| **Software** | **Version** | **Provider** | **Availability** |
|---|---|---|---|
| Python | 3.12 | Anaconda, Inc. | https://anaconda.org |
| Neuron | 7.7 |  | https://www.neuron.yale.edu/neuron |
| Simulation code |  | This paper | GitHub (link to be added) |

To evaluate the effects of TI stimulation across the hippocampal CA1 microcircuit, we simulated five neuronal classes: the Stratum Pyramidale Pyramidal Cell (PC), Parvalbumin-Positive Basket cell (PVBC), Axo-Axonic cell (AA), Ivy cell, and Oriens-Lacunosum Moleculare (OLM) cell.

Each model carries the electrophysiological profile assigned to it in the BBP dataset: cACpyr for the pyramidal cell, cNAC for PVBC and Ivy cells, bAC for the axo-axonic cell, and cAC for the OLM cell (Table 2). All cells were simulated at a physiological temperature of *T* = 37^*°*^C, with a fixed integration timestep of Δ*t* = min(0.025 ms, 1*/*20*f*_1_), where *f*_1_ is the carrier frequency of applied external electrical field. This ensure at least twenty steps for each carrier cycle.

**Table 2:** Neuronal models used in this study. Links provide access to the corresponding pages for more details and downloading the models.

| **Cell type** | **Cell layer** | **e-type** | **Cell id** |
|---|---|---|---|
| Pyramidal cell (PC) | SP | cACpyr | 1673 |
| Parvalbumin Basket cell (PVBC) | SP | cNAC | 408838 |
| Ivy cell (Ivy) | SP | cNAC | 432269 |
| Axo-axonic cell (AA) | SP | bAC | 430019 |
| Oriens Lacunosum Moleculare cell (OLM) | SO | cAC | 450290 |

Coordinates are those of the reconstructed morphology, with the soma at the origin. The soma-todendritic axis runs along *y*: in the pyramidal model the apical dendrites extend toward positive *y* (centroid +279 *µ*m, total extent 746 *µ*m) and the basal dendrites toward negative *y* (centroid −61 *µ*m), while the apical and basal centroids lie within 21 *µ*m of the soma along *x* and *z*. Field orientation is specified by the unit vector (*E*_*x*_, *E*_*y*_, *E*_*z*_) in this frame. The distal apical segment used throughout was located is 754 *µ*m from the soma by path distance.

### 4.2 Current injection

As a first approach to controlling the neuron’s operating point, we use NEURON’s *IClamp* point process to inject a constant current into either the soma or an apical dendrite segment. Because `IClamp` represents an electrode current, positive current depolarizes the membrane, thereby shifting the baseline membrane potential of the targeted segment. The threshold current *I*_thresh_ was determined independently for each model and each injection site by stepping 5% away from an initial estimate, growing the step up to a factor of two until the threshold was bracketed, then bisecting on a logarithmic scale to 0.1% precision. Threshold was defined as the lowest amplitude producing at least one spike, detected with textttscipy.signal.find_peaks using a height threshold of 0 mV at the soma (−30 mV in dendrites), a minimum prominence of 20 mV, and a minimum inter-peak interval of 7 ms. The injected current is reported throughout as *Ĩ* = *I/I*_thresh_, so that *Ĩ* = 1 corresponds to the threshold current. Because the conductance manipulations in Fig. 7 alter excitability, *I*_thresh_ was re-measured for each modified model.

### 4.3 Extracellular field stimulation

In the standard cable equation, the extracellular space is treated as an equipotential reference, so that the membrane potential is simply the intracellular potential, *V*_*m*_ = *V*_*i*_, and the cable equation is written only for the intracellular domain [64, 65]. Inserting NEURON’s `extracellular` mechanism adds two extracellular layers as explicit electrical domains, with the membrane potential defined relative to the perimembrane layer (*e*_0_),

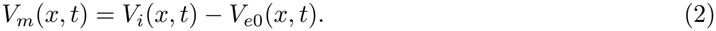

We left the extracellular layer parameters at their NEURON defaults (*x*_*raxial*_ = 10^9^ MΩ/cm, *x*_*g*_ = 10^9^ S/cm^2^, *x*_*c*_ = 0 *µ*F/cm^2^, two layers). At these values, the layers are effectively tied to ground, so *V*_*e*0_ follows the prescribed `e_extracellular` and the neuron does not influence its own extracellular potential. The field is therefore an imposed boundary condition rather than a quantity solved jointly with the membrane potential, and the system reduces to the usual cable equation with an extracellular driving term. This treatment neglects ephaptic effects, which is appropriate for an isolated neuron in a large conductive volume but would need revisiting in a densely packed population.

Following Aberra et al. [51], the field is applied through the `xtra` mechanism, which sets the extracellular potential at each segment according to its position:

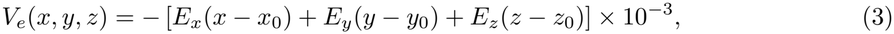

with coordinates in *µ*m and field components in V/m, giving *V*_*e*_ in mV. The origin (*x*_0_, *y*_0_, *z*_0_) was placed at the segment receiving the current injection, so that somatic and dendritic stimulation are compared symmetrically. Reported field amplitudes refer to the magnitude of **E** rather than to its individual components.

### 4.4 Simulating Temporal Interference

Following the methodology described in the later section, cells were exposed to a field of the form

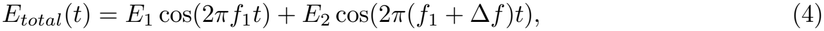

so that for Δ*f >* 0 the field is amplitude-modulated with envelope frequency Δ*f*, and for Δ*f* = 0 it reduces to an unmodulated sinusoid. This setup represents the field in the target region, where the interference is maximal.

The field was ramped on linearly over 100 ms, starting at half the simulation duration, and all analyses were restricted to the steady-state window following the ramp.

Modulated and unmodulated fields were matched by peak amplitude, following the original experimental demonstration, in which all conditions were applied at matched total current [2], and subsequent computational work [10, 11]. Writing each component of the modulated field as *A* cos(2*πft*), the cross term averages to zero over many carrier and beat cycles, so the modulated field has mean-square amplitude *A*^2^, whereas the unmodulated field of equal peak 2*A* has mean-square amplitude 2*A*^2^. Peak matching therefore gives the unmodulated condition twice the mean-square field. As a control, we repeated the comparison with the unmodulated field scaled by 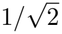, equalising the mean-square amplitude of the two conditions (see Appendix).

For the orientation analysis (Fig. 4), field direction was sampled at 100 angles uniformly spaced over 2*π* (3.6^*°*^ resolution) within each of the three orthogonal planes.

### 4.5 Simulating Synaptic Inputs

To examine neuronal responses under more physiologically relevant conditions, we incorporated excitatory synaptic inputs distributed across the somatodendritic arbor of the CA1 pyramidal neuron model. Synaptic inputs were modeled using the `Exp2Syn` mechanism in NEURON, which implements a dual-exponential conductance waveform:

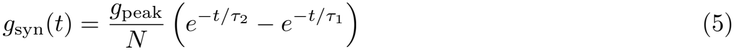

where *τ*_1_ and *τ*_2_ are the rise and decay time constants, respectively, and *g*_peak_ is the peak synaptic conductance. The normalization constant

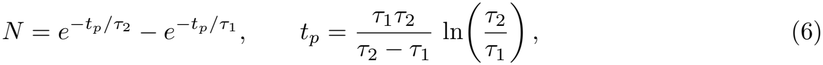

ensures that the waveform attains its maximum *g*_peak_ at *t* = *t*_*p*_, as implemented in NEURON [63]; for the values used here, *t*_*p*_ = 21.5 ms and *N* = 0.582. The synaptic current is then given by

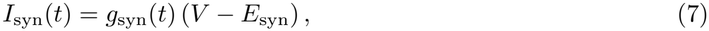

where *E*_syn_ is the synaptic reversal potential.

Excitatory synapses were modeled as NMDA-type conductances (*τ*_1_ = 10 ms, *τ*_2_ = 60 ms, *E*_syn_ = 0 mV), consistent with the slow, sustained depolarization characteristic of NMDA receptor activation. This implementation provides slow excitatory conductances but no voltage-dependent magnesium block, and no inhibitory input was included; it is intended to reproduce the conductance load and voltage fluctuations associated with ongoing excitatory drive rather than the full synaptic complement of a CA1 pyramidal neuron. Each synapse received input from an independent `NetStim` presynaptic source firing at a mean rate of 50 Hz with a noise level of 0.5 (interpolating between periodic and fully Poisson spiking), and a unitary peak conductance of 30, 35 or 40 pS. Synapses were placed on a random subset of segments across all somatodendritic compartments, excluding the axon, with the fraction of targeted segments controlled by a ratio parameter (ratio = 0.1 unless otherwise stated). The model comprises 132 somatodendritic segments, each independently selected with probability 0.1, giving 13.2 ± 3.4 synapses (mean ± SD) and total synaptic conductances of approximately 0.40, 0.46 and 0.53 nS respectively, with the subset redrawn for each simulation trial. Each condition was simulated for 10 s across 100 independent trials, differing in the synaptic subset, the presynaptic spike trains, and the random seed; the first 5 s were discarded and firing rates computed over the remaining 5 s.

This configuration produces sustained depolarization and membrane noise arising from the synaptic dynamics rather than imposed externally, providing a parametric axis from weakly to strongly driven conditions. The total synaptic conductance is nonetheless modest relative to the high-conductance states reported in vivo [39], so these results characterize the onset of conductance-dependent attenuation rather than its asymptotic limit.

### 4.6 Quantifying Resonance from Voltage and Current Responses

Classical approaches to characterizing neuronal resonance inject a sinusoidal current of fixed amplitude and swept frequency, a chirp or ZAP (impedance amplitude profile) stimulus, and compute the membrane impedance as the ratio of the Fourier transform of the voltage response to that of the injected current [66, 14, 17, 41]. The frequency at which this impedance is maximal defines the resonant frequency, and the sharpness of the peak quantifies the resonance strength (Q-factor) [14].

In the present study, the electric field serves as the oscillatory input rather than an injected current, and the relevant quantity is how strongly the membrane responds to fields of different envelope frequencies. Under a fixed-amplitude sinusoidal or amplitude-modulated field, the standard deviation of the somatic membrane voltage *σ*(*v*_*s*_) provides a direct measure of the amplitude of the field-driven voltage response. The frequency at which *σ*(*v*_*s*_) reaches its maximum was taken as the resonant frequency, providing a measure of frequency-selective membrane responsiveness without requiring explicit calculation of the impedance [67].

We additionally quantified *σ*(*I*_*Na*_), the standard deviation of the somatic sodium current, as a complementary measure of ionic channel recruitment at each envelope frequency. While *σ*(*v*_*s*_) reflects the aggregate membrane response, *σ*(*I*_*Na*_) isolates the contribution of the fast inward sodium current and provides a direct readout of how channel activation varies with envelope frequency. Both quantities were computed over a steady-state window following the onset of the field, excluding transients. Where the time-averaged sodium current ⟨*I*_*Na*_⟩ is reported instead, it measures net sodium influx over the analysis window, as distinct from *σ*(*I*_*Na*_), which measures the amplitude of its field-driven modulation. Sodium current is plotted as influx throughout, with inward current taken as positive.

*I*_K_ denotes the total potassium current at the soma, recorded as NEURON’s *ik* variable, which sums the contributions of all potassium mechanisms present in the model; the individual mechanism currents, including *I*_Kdr_ and *I*_M_, were recorded separately.

Under synaptic drive, where the membrane potential fluctuates spontaneously, the field-evoked change in firing was quantified as the difference in mean firing rate between field-on and field-off conditions. The difference rather than the ratio was used because baseline firing approached zero at the lowest synaptic conductance tested, which renders ratio measures unstable.

## 5 Acknowledgements

R.P.A., L.L.G., S.C. and C.R.M. acknowledge funding from the Fundación Instituto Zapata Briceño de Neurociencia para la Investigación de la Inteligencia Humana. R.P.A., L.L.G. and C.R.M. acknowledge funding from the Ministerio de Ciencia, Innovación y Universidades and Agencia Estatal de Investigación through the grant CEX2021-001164-M. R.P.A., S.C. and C.R.M. were supported by grant PID2024-162400OB-C21/-C22. S.C. was supported by grant CEX2021-001165-S, MICIU/AEI/10.13039/501100011033 and FEDER, UE. L.L.G. was supported by the project ANCHOR (PID2024-162343OB-I00). F.C. and G.R. have received funding from the European Research Council (ERC) under the European Union’s Horizon 2020 research and innovation programme (grant agreement No 855109, GALVANI).

## Appendix

### Local subthreshold resonance along the apical trunk

To better understand the differences in the resonant properties between the soma and the apical segment (Fig. 1), we sampled 10 apical segments evenly spaced in path distance along the path from the soma to the most distal apical segment of the CA1 pyramidal cell, as shown in Fig. 8. At each location, the threshold was found using the methodology described in 4.2. The model was simulated for 10s, where the first 7.5s where discarted and the 2.5s used for the threshold estimation. At 341 *µ*m, dendritic spikes preceded somatic spikes by 0.85 ms, confirming local initiation.

We used three protocols, each answering a different question. In all of them, every location was held by a DC current at a fraction of its own threshold and driven sinusoidally.

- *Field, fixed clamp*: Each location was held at 95% of its threshold and exposed to a uniform field (*E*_0_ = 1 V/m), applied as an extracellular potential as described in 4.4. This gives the field-driven resonance with every location at the same fraction of its threshold.
- *Field, scaled clamp*. A uniform field polarizes the membrane very unevenly along the trunk: strongly at the soma and in the tuft, weakly in the mid-trunk (see Fig. 8c). At a fixed holding fraction, weakly polarized locations therefore stay far from threshold relative to what the field can do there. In this protocol the field was unchanged, and each location’s holding current (90– 99.5% of threshold) was set so that the field’s peak response covered the same fraction of its remaining distance to threshold. Adjusting the clamp rather than the field keeps the drive the same for the whole cell.
- *Current, fixed clamp*. To isolate each location’s intrinsic resonance from the way the field couples to the cell, the field was replaced by a sinusoidal current injected at the same segment, on top of a holding current at 95% of threshold. Its amplitude, 0.5% of threshold, was the largest tested that kept every location subthreshold; larger amplitudes evoked spiking at the distal tip even though their peak current stayed below the DC threshold.

The field was applied as an extracellular potential, as described in 4.4, with *E*_amp_ = 1.0 V/m, and responses are reported per unit *E*_0_. In the scaled-clamp protocol, each location’s holding fraction *α* was set so that the field’s peak response covers half of the remaining distance to threshold, with that distance expressed as a voltage through *I*_*th*_*R*_*in*_:

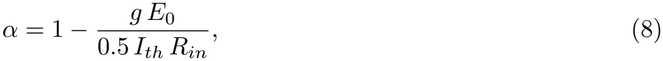

where *g* is the location’s largest subthreshold response per V/m in the field, *Ĩ* = 0.95 protocol. *α* was bounded to 0.90–0.99 (0.995 at 173 *µ*m) and ranged from 0.90 at the tip to 0.995 in the mid-trunk. Near threshold, the intrinsic local resonance of the apical trunk, measured with a local sinusoidal current, is about 11–12 Hz from the soma to roughly 500 *µ*m and rises steeply to about 30 Hz in the distal tuft (Fig. 8b, circles). The field-driven resonance follows this profile where the field strongly polarizes the membrane, at the soma and in the distal tuft. In the mid-trunk, where the polarization by a uniform field changes sign and the response is weakest (Fig. 8c). Holding these locations closer to threshold compensates for this effect.

### Resonance across cells

Figure 9 summarizes the somatic resonance frequencies of the cells studied in this work. Following the protocols described in the later section, a DC current clamp (IClamp) was placed at the soma and set to *Ĩ* = 0.95. A uniform electric field of amplitude 2 V/m was then applied, with a pure sinusoidal waveform. Each panel shows the response, measured as the standard deviation of the membrane voltage at the soma.

**Figure 9:**
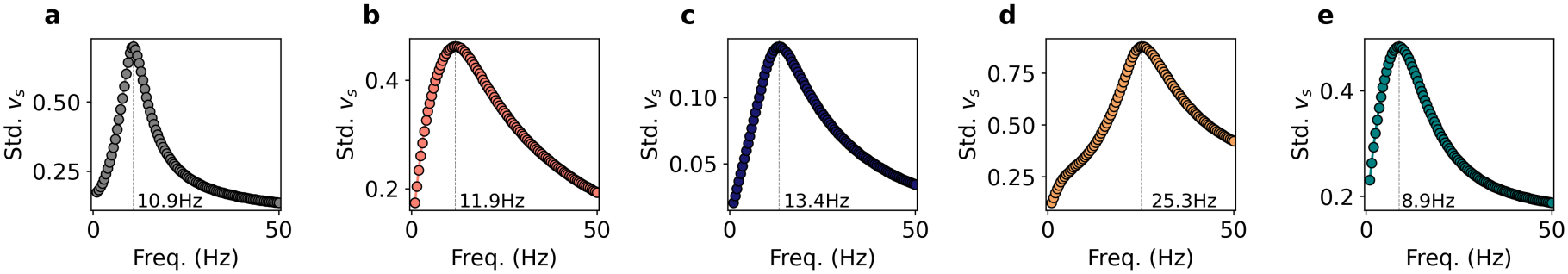
Somatic membrane voltage resonance across stimulation frequencies. Standard deviation of somatic membrane voltage as a function of stimulation frequency at 99.5% of the threshold current and an electric field amplitude of 2 V/m. Resonant profiles are shown for (a) PC, (b) PVBC, (c) AA, (d) Ivy, and (e) OLM cells.

We note that resonant frequencies reported for CA1 pyramidal neurons vary across studies, and the experimental protocols differ in several respects that could contribute to this. Narayanan and Johnston [41] recorded from young adult rats (5–7 weeks), prepared with transcardial perfusion of ice-cold cutting solution before slicing, and do not state a recording temperature. Zemankovics et al. [18] used juvenile animals (postnatal day 14–26), decapitated under isoflurane without perfusion, with recordings made at 34–37 °C. Hu et al. [44] recorded at 32.9 ± 0.09 °C and were the only group to block synaptic transmission pharmacologically, adding DNQX and bicuculline to the extracellular medium, with TTX and a low-calcium solution for the voltage-clamp measurements of *I*_*M*_ and *I*_*h*_. The three protocols therefore differ in animal age, recording temperature, and whether the measured resonance includes a synaptic contribution. We do not attempt to attribute the reported differences to any one of these factors, but note that comparisons of resonant frequency across studies, and between experiment and model, should take them into account. Of the three, the preparation of Hu et al. is closest to the conditions simulated here, in which synaptic input is absent, and the resonance is intrinsic to the membrane.

### Field scaling

To understand the band in Fig. 3 where the modulated field drives spiking while the unmodulated field does not, we examined how the subthreshold response scales with field amplitude. At a fixed normalized input current (*Ĩ* = 0.98), we measured the standard deviation of the somatic membrane potential and of the K^+^ and Na^+^ currents as a function of field amplitude, for each carrier frequency and both waveforms (Fig. 10), and fitted each curve with a power law (Fig. 11). The unmodulated response grows linearly with field amplitude at every carrier, whereas the modulated response grows supralinearly, increasingly so at higher carriers. A supralinear response reaches the spiking threshold over a shorter range of field amplitudes, which is consistent with the modulated field recruiting firing where the unmodulated field does not, and with the shape of this region changing with the carrier. The power-law model is given by

**Figure 10:**
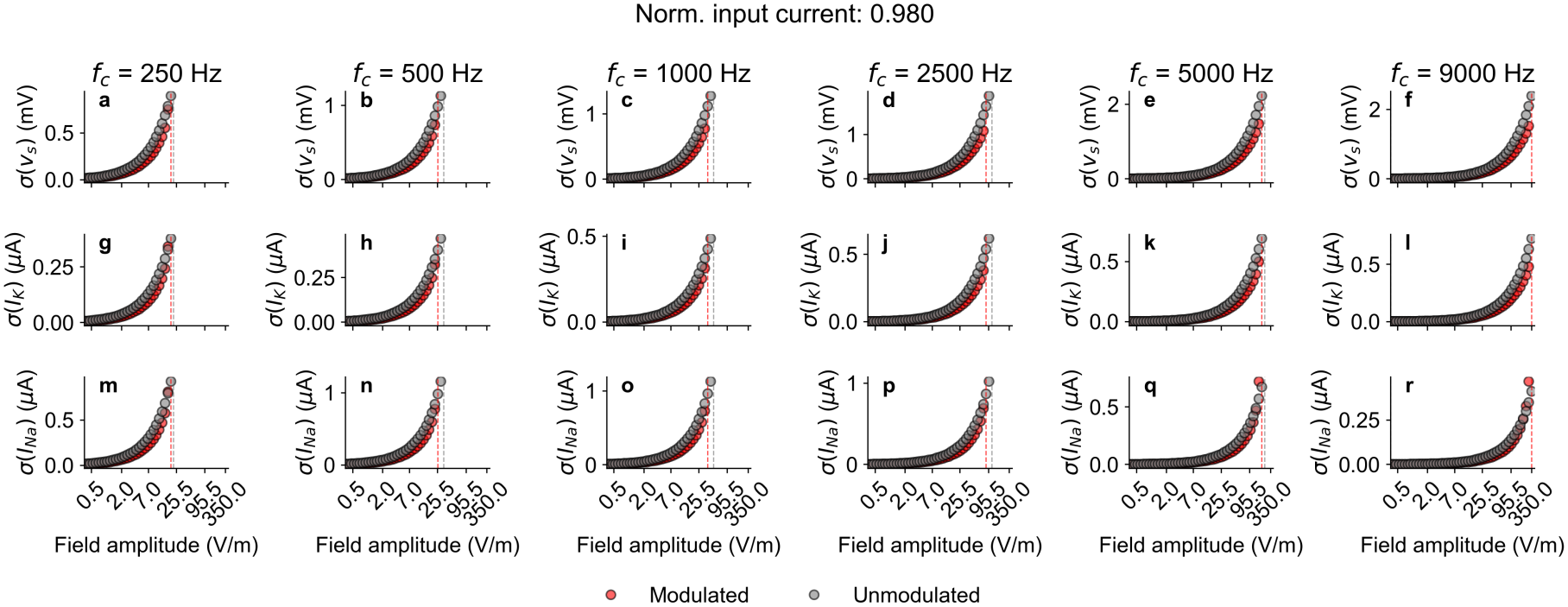
Sames as figure 3 but with fixed normalized current input. First row shows the standard deviation of the somatic membrane voltage, second row the std. of K current and third the std. of the Na current.

**Figure 11:**
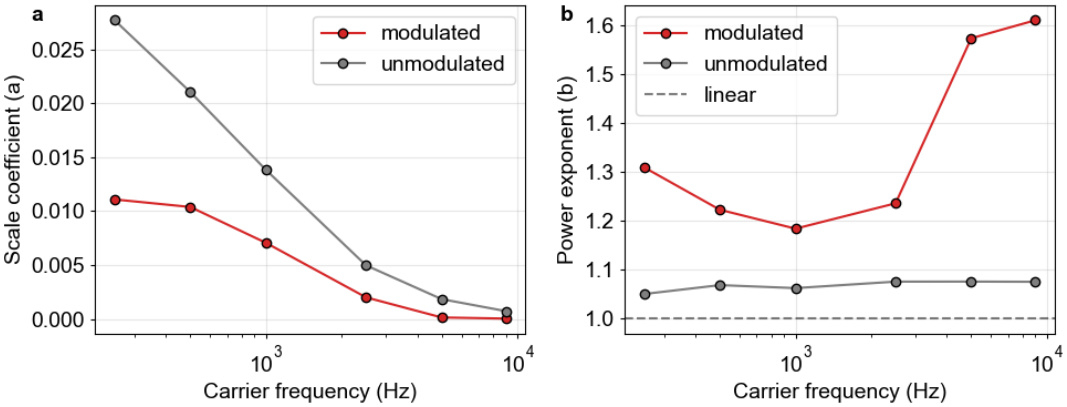
From figure 8 we estimate how the response to the field scales with the carrier frequency (for the same normalized input current).

**Figure 12:**
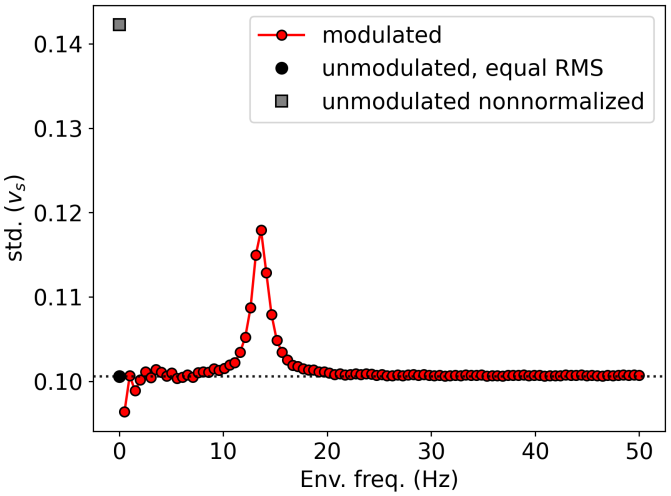
Equal-RMS control in the axo-axonic cell. Standard deviation of the somatic membrane potential as a function of envelope frequency (red), with the unmodulated reference shown at matched peak amplitude (grey circle) and at matched mean-square amplitude (black circle); the dotted line marks the latter.

**Figure 13:**
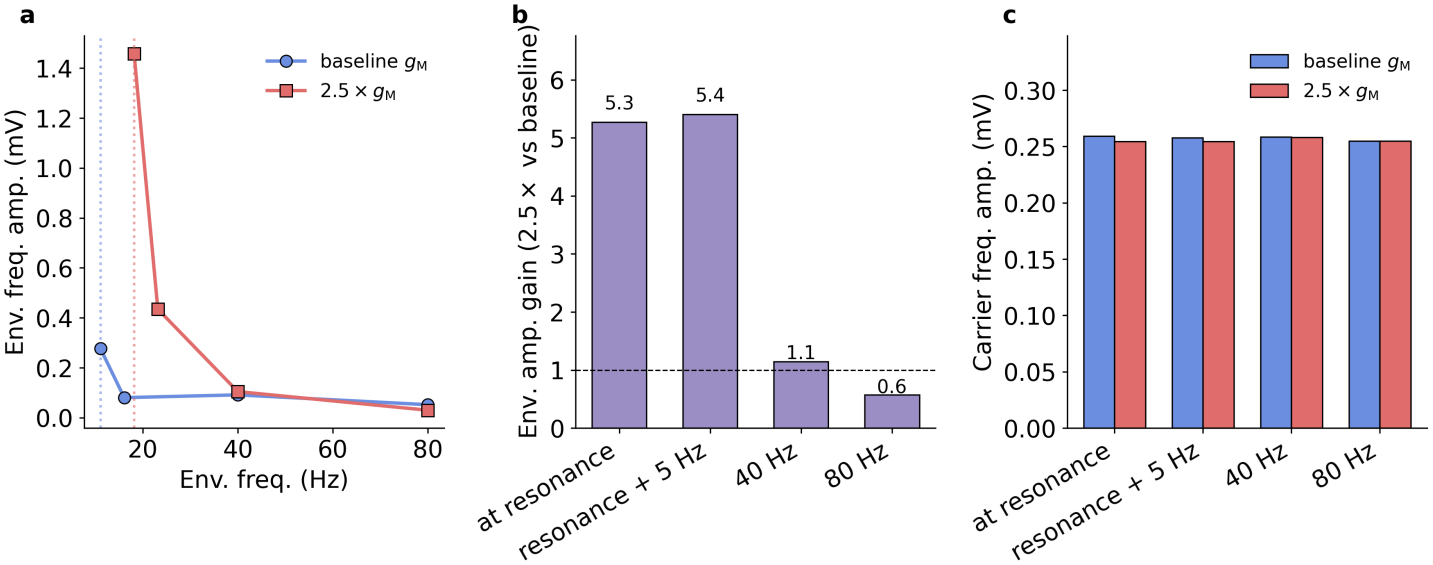
Increasing 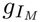 enhances amplification, not demodulation, in the PVBC cell. a, Amplitude of the beat-frequency component of the somatic membrane potential as a function of beat frequency, at baseline 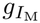 (blue) and at 2.5*×* baseline (red). Dotted vertical lines mark each condition’s own resonant frequency, measured from its direct sinusoidal response. b, Ratio of the two conditions at each beat frequency. The enhancement is confined to the resonant band, reaching 5.3*×* at each cell’s own resonance and 5.4*×* at resonance plus 5 Hz, and is absent at 40 and 80 Hz. c, Amplitude of the carrier-frequency component, which is unchanged by the manipulation across all conditions.

**Figure 14:**
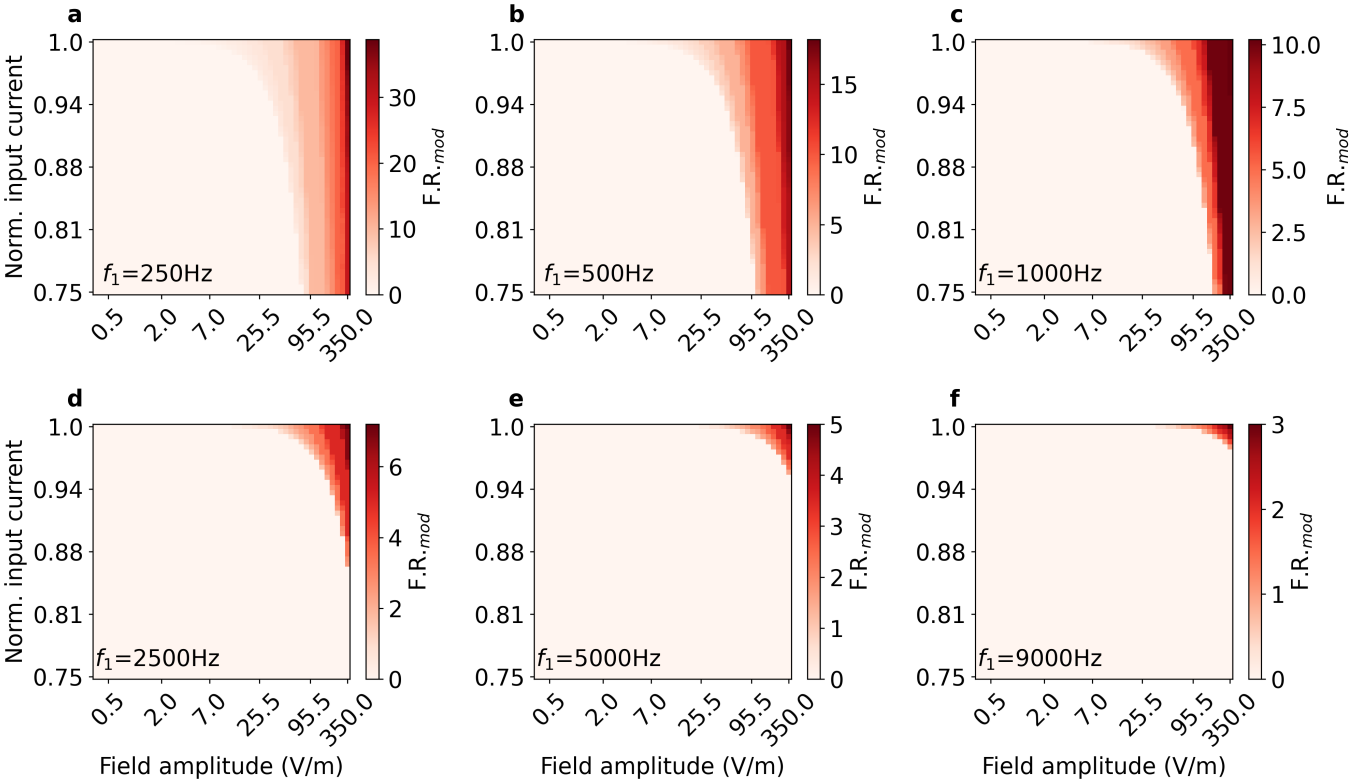
Firing-rate maps of Fig. 3 for the modulated field.

**Figure 15:**
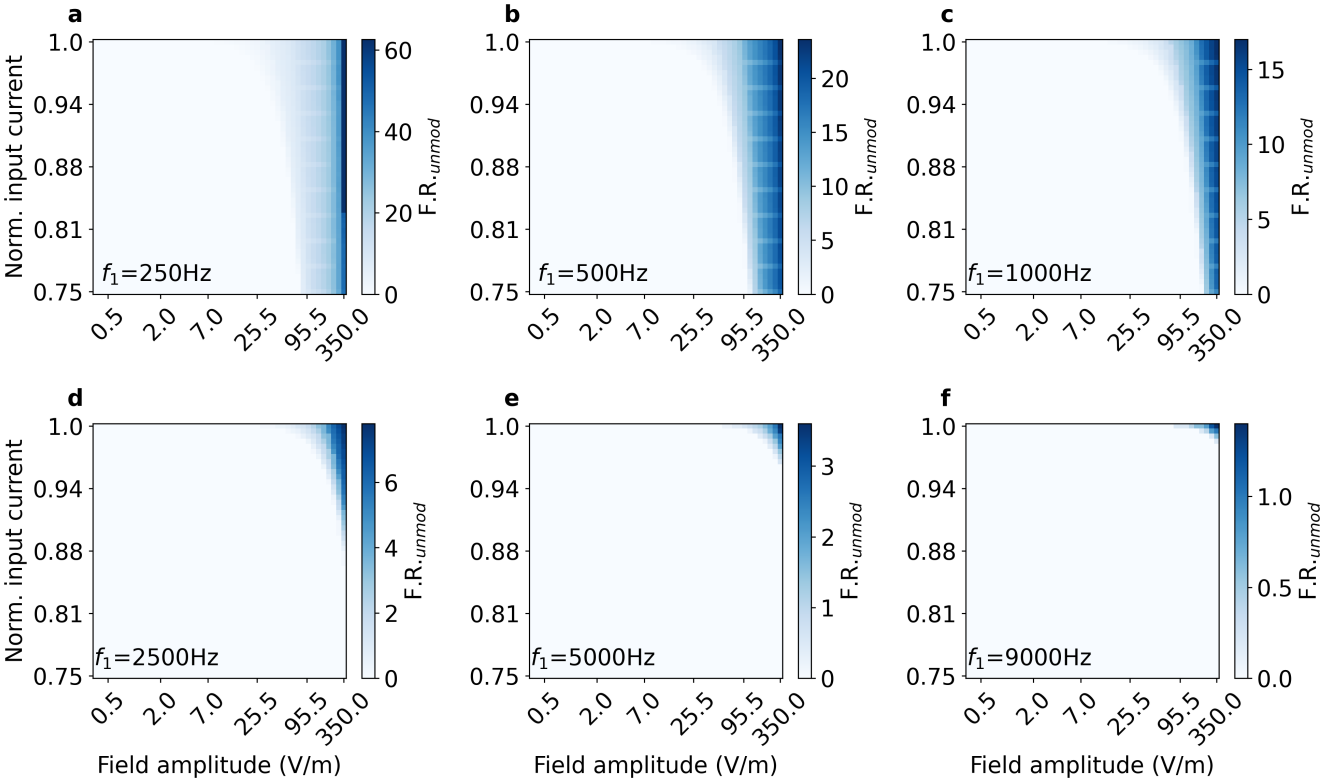
Firing-rate maps of Fig. 3 for the unmodulated field.

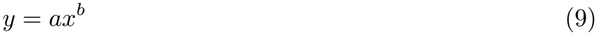

where *a* is the scale coefficient and *b* is the power exponent.

Such a model accurately described the input–output relationship in both conditions, with consistently better fits for the unmodulated waveform (*R*^2^ = 0.9998 *±* 0.0001, residual = 0.00014 *±* 0.00004) than for the modulated waveform (*R*^2^ = 0.9843 *±* 0.0065, residual = 0.00845 *±* 0.00525). In the unmodulated condition, the exponent remained close to unity at every carrier frequency (*b* = 1.035 *±* 0.008), indicating an approximately linear scaling of the response with field amplitude. The modulated condition was supralinear throughout (*b* = 1.19–1.63), with the exponent close to 1.2–1.3 for carriers up to 2500 Hz and rising to ≈1.6 at 5000 and 9000 Hz. The scale coefficient decreased with carrier frequency in both conditions, as expected from low-pass attenuation of the carrier, but only the modulated waveform showed a concomitant increase in the exponent. Raising the carrier under amplitude modulation therefore does not simply reduce the overall response magnitude; it shifts the input–output relationship toward stronger supralinear scaling.

### Equal RMS comparison

Modulated and unmodulated conditions are matched by peak amplitude throughout the main text, following the convention of the experimental and modelling literature. At matched peak the unmodulated field carries twice the mean-square amplitude of the modulated one, since the cross term of the two detuned components averages to zero over many beat cycles. To isolate the contribution of the waveform from that of delivered energy, we repeated the comparison with the unmodulated field scaled by 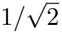.

### Effects of I_*h*_ manipulation on demodulation and resonant amplification

A TI response at the beat frequency requires two steps: a nonlinearity must generate a component at that frequency, since the applied field contains none, and the membrane must then amplify it. Increasing a resonant conductance could in principle act on either. The two make different predictions: a change in generation would scale the response at all beat frequencies comparably and might alter the carrier response, whereas a change in gain would act selectively near the resonant peak and leave the carrier unaffected. To distinguish them, we measured the Fourier component of the somatic membrane potential at the beat frequency, rather than its total standard deviation, at beat frequencies spanning the resonant peak and well beyond it.

